# Vertical distribution and migration of microplastics in soils from Fars Province, southwest Iran

**DOI:** 10.1101/2025.09.17.676919

**Authors:** Shekoufeh Forouzan, Sajjad Abbasi, Ali Akbar Moosavi, Majid Baghernejad, Sayyed Mahmoud Enjavinezhad, Andrew Turner

## Abstract

Microplastics (MPs) in soils have recently emerged as a significant environmental concern because of their potential impacts on ecosystems and human health. In this study, MPs have been determined in genetic soil horizons to a maximum depth of 140 cm along four transects that encompass various land uses (managed and unmanaged) in Fars Province, southwest Iran. Soils have also been characterised in terms of texture and chemistry using established methods. With little contemporary or historical application through agricultural practices, MPs were dominated by fibres of various colours and sizes and polymeric construction (mainly polyamides, polyesters and polyolefins), with remaining particles largely consisting of sheet-like fragments. MP abundance (up to about 200 per kg of dry soil) and size were heterogeneously distributed throughout the region and with respect to soil depth, regardless of land use, with inverse correlations with soil particle size observed at two locations. We infer that atmospheric deposition is the principal source at the soil surface and that MPs that evade erosion are able to readily migrate downwards to depths extending to at least that of the lowest horizon sampled. Migration appears to be independent of particle size or density and is likely driven by percolating precipitation but facilitated through bioturbation and soil cracking during dry periods. The persistence and vertical migration of MPs in soils may have adverse impacts on subterranean ecosystems and ground water quality.

**Highlights:**

- Microplastics (MPs) determined in soil horizons covering different land uses in Fars Province
- MPs dominated by fibres and derived largely from atmospheric deposition
- MPs distributed heterogeneously with depth, land use, soil texture
- Regardless of size and density, MPs able to migrate to at least 140 cm
- Migration may have adverse impacts on subterranean ecosystems and ground water

## 1. Introduction

Despite the majority of microplastic (MP) sources being land-based, studies in the terrestrial environment have emerged relatively recently (Horton et al., 2017). Soils receive MPs through atmospheric fallout (Klein and Fischer, 2019), practices related to land use (Cusworth et al., 2024), and contamination from local industrial or urban sources (Afrin et al., 2020). Soils can then act as a secondary source of MPs to the atmosphere through surface erosion (Abbasi et al., 2023) or a receptor that has potential impacts on communities and habitats or soil structure, aeration, fertility and nutrient cycling (de Souza Machado et al., 2019; Gao et al., 2021). Over time, MPs can also migrate downwards through soil with precipitation or via bioturbation and have the potential to impact groundwater resources (Ren et al., 2021).

Several studies have recently compared the effects of land use on the abundance and characteristics of MPs in soils. In general, these have focussed on different agricultural practices, including land managed for arable crops, pasture, orchards, plantations, greenhouses and forests (Álvarez-Lopeztello et al., 2021; Liu et al., 2022; Zhang et al., 2022a; Bi et al., 2023; He et al., 2023; Li et al., 2023; Qiu et al., 2023). In most cases, differences reported are related to soil characteristics (e.g., pH, texture, organic matter, nitrogen content), climatic factors (e.g., precipitation, temperature, wind speed) and/or practices that involve the direct introduction of plastics from mulching, fertilisers, sludge or irrigation water. However, differences are location- or activity-specific and are not always statistically verified, and data are usually reported for surface soils or samples down to typical tillage depth (about 25 cm; Hobson et al., 2022). This means that a general understanding and prediction of the key drivers for MP distributions in soils and their propensity for migration is lacking.

In the present study, we investigate the abundance and characteristics of MPs in soils from an arid-semi humid region of Iran (Fars Province) that encompasses various managed and unmanaged land uses, including barren land where degradation takes place under wind erosion, but where distinct sources of MPs are lacking. We examine genetic soil horizons to depths exceeding one metre and determine key physical and chemical soil properties with the aim of improving our broader understanding of the vertical distribution and persistence of MPs and their potential for impacting on subterranean ecosystems and groundwater.

## 2. Material and methods

### 2.1. Study area

Fars province is located in southwest Iran. It covers an area of about 120,000 km^2^ and is home to about 5 million people, with about a third of the population concentrated in the provincial capital city of Shiraz. The more mountainous, northern areas of Fars have cold winters and mild summers, while southern Fars experiences cold winters but hot summers; the central region has mild and relatively wet winters and hot, dry summers (Haghighi and Keshtkaran, 2014). The average annual precipitation for the whole province is 330 mm.

The agricultural sector of Fars plays a key role in the production and security of food for Iran and makes a significant contribution to the gross national product. It is also responsible for around 95% of total annual water consumption of about 10 billion m^3^ (Haghighi and Keshtkaran, 2014). Important agricultural products across the province include wheat, corn, rice, dates, barley, figs, pomegranates, cotton, walnuts, pistachios, citrus fruits, sugar beets and tomatoes.

#### 2.2. Sampling

Sampling was undertaken from the calcareous soils of the alluvial plains of four regions: Dasht Arjan, Darab, Sarvestan and Sepidan (Figure 1; Table 1); that encompassed different land uses. Specifically, agriculture for wheat and pistachios, second and third grade pastures (low vegetation cover), oak forest, private gardens and barren land subject to erosion and with some evidence of historical agricultural practices. In each region, six to eight profiles (up to 1.4 m in depth) along a sloping transect of between about 3 and 10 km were exposed by digging with a stainless-steel shovel. Approximately 2 kg of soil (*n* = 73 in total), collected from each visible, genetic horizon (two to four) with a smaller shovel, were stored in aluminium foil and transported to the laboratory where they were passed through a 2-mm stainless-steel sieve.

**Table 1:**
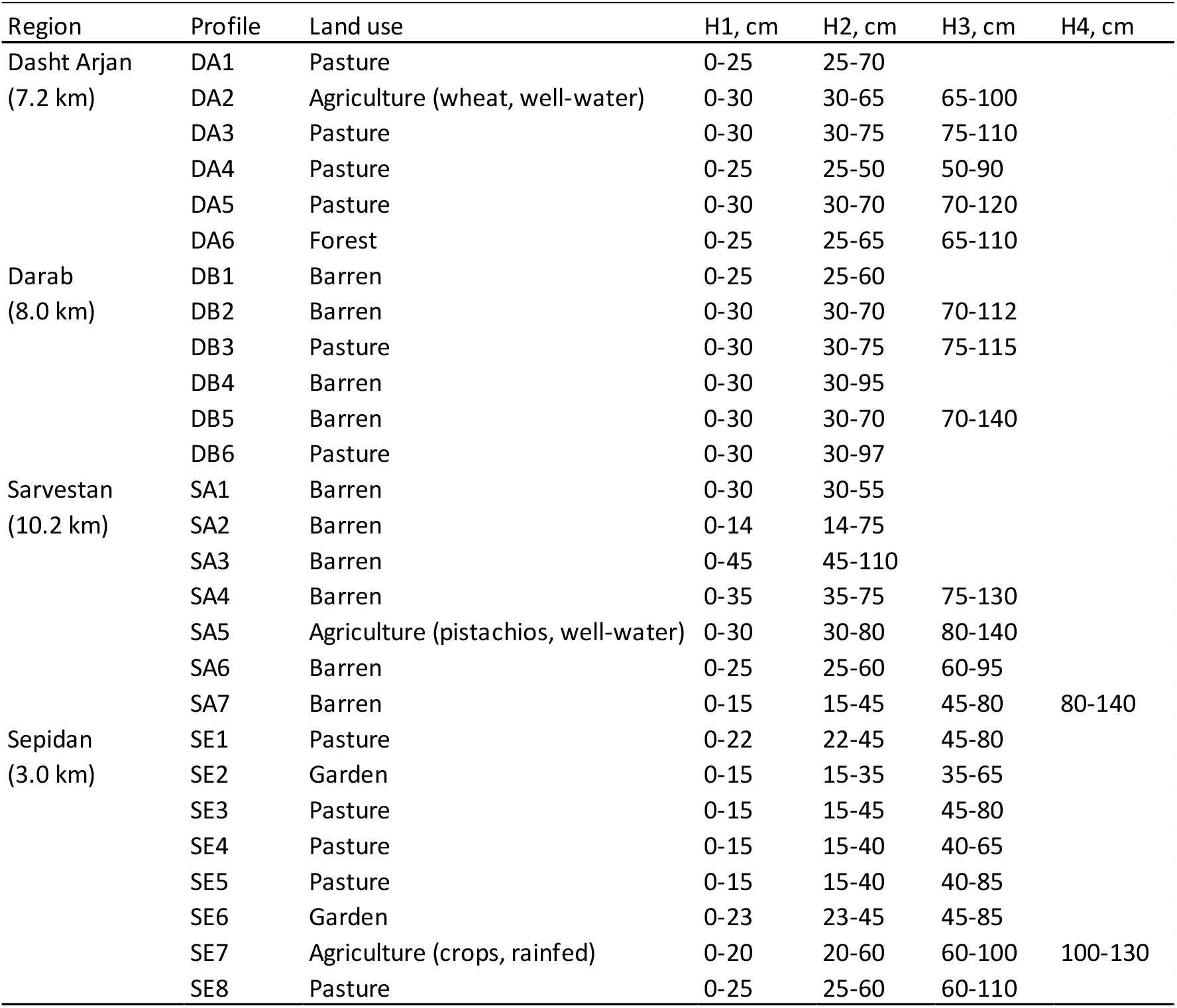
Land uses and profiles and genetic horizons sampled from the four regions of Fars Province under study. In parentheses are the lengths of each transect. Note, for agriculture the crops grown at the time of sampling are shown along with the source of irrigation water.

| Region | Profile | Land use | H1, cm | H2, cm | H3, cm | H4, cm |
| --- | --- | --- | --- | --- | --- | --- |
| Dasht Arjan<br>(7.2 km) | DA1 | Pasture | 0-25 | 25-70 |  |  |
|  | DA2 | Agriculture (wheat, well-water) | 0-30 | 30-65 | 65-100 |  |
|  | DA3 | Pasture | 0-30 | 30-75 | 75-110 |  |
|  | DA4 | Pasture | 0-25 | 25-50 | 50-90 |  |
|  | DA5 | Pasture | 0-30 | 30-70 | 70-120 |  |
|  | DA6 | Forest | 0-25 | 25-65 | 65-110 |  |
| Darab<br>(8.0 km) | DB1 | Barren | 0-25 | 25-60 |  |  |
|  | DB2 | Barren | 0-30 | 30-70 | 70-112 |  |
|  | DB3 | Pasture | 0-30 | 30-75 | 75-115 |  |
|  | DB4 | Barren | 0-30 | 30-95 |  |  |
|  | DB5 | Barren | 0-30 | 30-70 | 70-140 |  |
|  | DB6 | Pasture | 0-30 | 30-97 |  |  |
| Sarvestan<br>(10.2 km) | SA1 | Barren | 0-30 | 30-55 |  |  |
|  | SA2 | Barren | 0-14 | 14-75 |  |  |
|  | SA3 | Barren | 0-45 | 45-110 |  |  |
|  | SA4 | Barren | 0-35 | 35-75 | 75-130 |  |
|  | SA5 | Agriculture (pistachios, well-water) | 0-30 | 30-80 | 80-140 |  |
|  | SA6 | Barren | 0-25 | 25-60 | 60-95 |  |
|  | SA7 | Barren | 0-15 | 15-45 | 45-80 | 80-140 |
| Sepidan<br>(3.0 km) | SE1 | Pasture | 0-22 | 22-45 | 45-80 |  |
|  | SE2 | Garden | 0-15 | 15-35 | 35-65 |  |
|  | SE3 | Pasture | 0-15 | 15-45 | 45-80 |  |
|  | SE4 | Pasture | 0-15 | 15-40 | 40-65 |  |
|  | SE5 | Pasture | 0-15 | 15-40 | 40-85 |  |
|  | SE6 | Garden | 0-23 | 23-45 | 45-85 |  |
|  | SE7 | Agriculture (crops, rainfed) | 0-20 | 20-60 | 60-100 | 100-130 |
|  | SE8 | Pasture | 0-25 | 25-60 | 60-110 |  |

**Figure 1.**
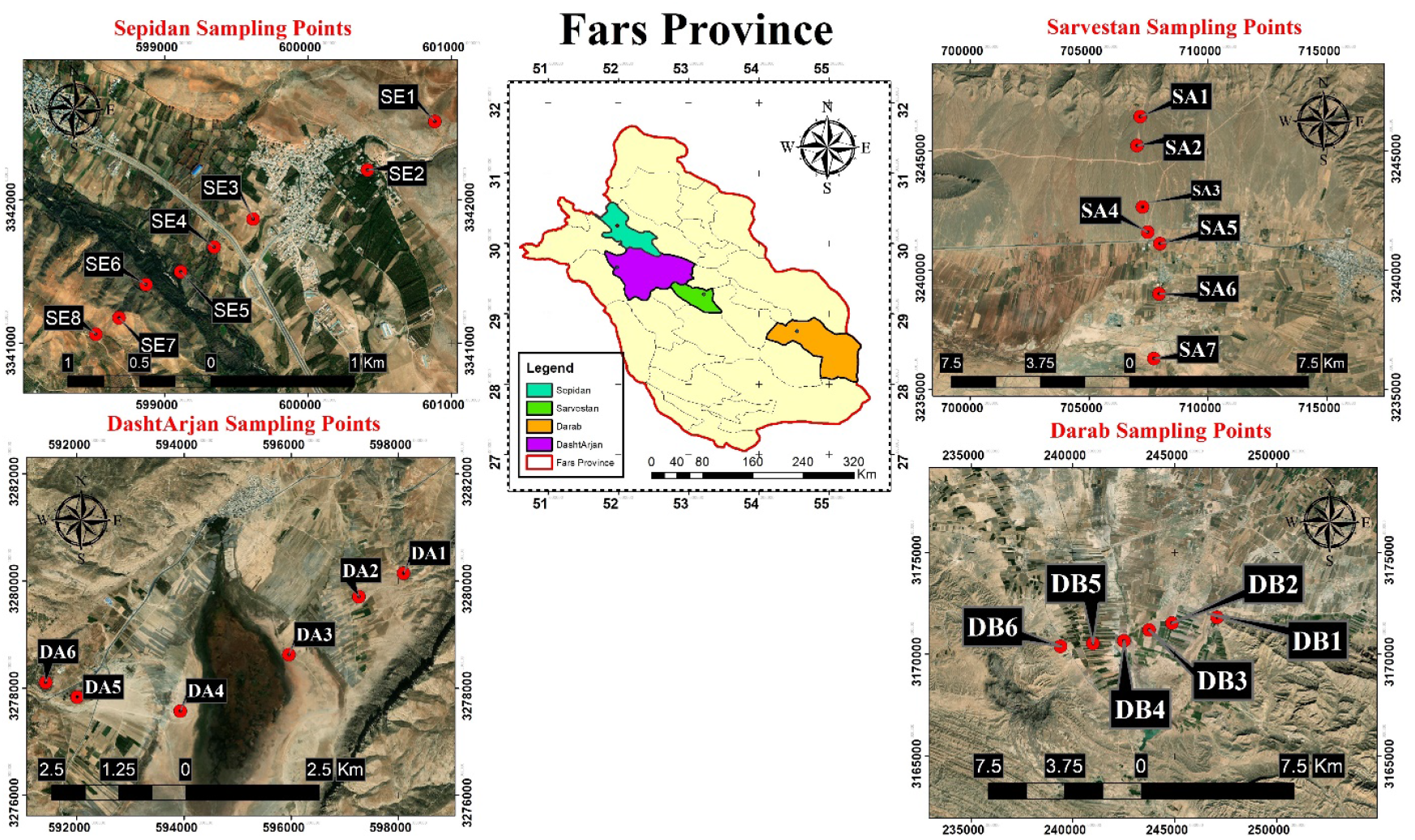
Location of the four transects and soil profiles sampled in Fars Province (The original maps were extracted from https://earthexplorer.usgs.gov and some needed modifications in Arc GIS 10.8 were performed by the authors).

#### 2.3. Microplastic separation

Fifty g of each soil sample was transferred to a cleaned (washed in 50% HCl and rinsed with deionised water) 600 mL glass beaker using a stainless-steel spoon before the contents were loosely covered with foil and air-dried in a laminar flow hood (without airflow) for 24 h at 25 °C.

In new (and cleaned) 600 mL glass beakers, organic matter was decomposed by oxidising 50 g of each sample in 300 mL of 30% H_2_O_2_ solution (Arman Sina, Tehran) at 25°C until the cessation of bubble formation. The remaining material (and H_2_O_2_) was washed through a 150-mm diameter S&S filter paper (blue ribbon cellulose circle, grade 589/3, 2-μm pore size) with filtered, deionised water before residues, on their filters, were dried for 2 h in a sand bath at 60°C and stored in individual glass Petri dishes.

Dried residues were added to saturated, 300 mL solutions of ZnCl_2_ (Arman Sina, Tehran; density ∼ 1.7 g cm^-3^) in a series of cleaned, 600 mL glass beakers. Following a period of initial agitation at 350 rpm on a lateral shaker, the contents were allowed to settle for 24 h before the decanted contents were vacuum-filtered through S&S filter papers. This procedure was repeated twice, with the resulting filters being air-dried for 72 h at 25°C in the laminar flow hood (without airflow) and transferred to individual Petri dishes.

#### 2.4. Microplastic identification and characterisation

Filters were inspected under a binocular microscope (Carl-Zeiss) at magnifications of up to 200x, aided by ImageJ software, to identify, quantify and characterise microplastics. Criteria for identification included thickness, cross-sectional properties, shininess, hardness, surface structure and their response to a hot, 250 μm stainless steel probe. Size measurements were based on the length along the longest axis (*L*) and particles were classified as follows: *L* ≤ 100 µm (with a detection limit of about 30 μm), 100 < *L* ≤ 500 µm, and *L* ≥ 500 µm. Colour was categorised as white-transparent, yellow-orange, red-pink, blue-green or black-grey, and shape was categorised as fibre (with a length to diameter ratio of > 3), film-fragment or spherule-granule.

The polymeric makeup of 98 MPs of a range of shapes, sizes and colours from different locations (specifically, from profiles DA5, DB1, DB3, DB4, DB5, SA1, SA3 and SE7) was determined using a micro-Raman spectrometer (µ-Raman-532-Ci, Avantes, Apeldoorn, Netherland) with a laser of 785 nm and Raman shift of 400-1800 cm^-1^.

### 2.5. Minimisation and assessment of microplastic contamination

In the laboratory, windows and doors remained closed, benches were thoroughly cleaned using ethanol and attire was exclusively composed of cotton-based materials. All reagents and solutions employed (including deionised water) were vacuum-filtered through 2 μm before being used and aluminium foil was used to cover equipment while not in use.

Contamination in the laboratory was assessed by processing ten aliquots of filtered, deionised water according to the protocols outlined above. Microscopic analysis of these filters revealed no detectable MPs.

#### 2.6. Physical and chemical properties of soils

Physical and chemical parameters of the soils or their saturated pastes or extracts (in pure water) were measured by established techniques (e.g., Puffeles and Nessim, 1957; Gee and Bauder, 1986; Ryan et al., 2007). Briefly, organic carbon was measured by wet oxidation with potassium dichromate and sulphuric acid; texture was determined by hydrometry with sodium hexametaphosphate as a dispersing agent; pH and electrical conductivity were measured directly in pastes and extracts, respectively; soluble 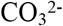 and 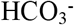 concentrations were determined by titration with sulphuric acid; calcium carbonate equivalent was determined by neutralisation with HCl; soluble Mg and Ca concentrations were determined by titration with EDTA; and soluble Na and K concentrations were measured directly in extracts using a Corning 405 flame photometer.

#### 2.7. Data presentation and statistics

MP data are presented as both number per sample (50 g) or concentration on a dry weight basis (number per kg). Parametric or non-parametric correlations and difference testing between variables were undertaken for each region using SPSS 26.0 (IBM) after checking for data normality using a Kolmogorov-Smirnov test, with an α-value for significance of 0.05.

### 3. Results

#### 3.1. Soil characteristics

The physical and chemical characteristics of the soils from each transect (region), profile and genetic horizon are summarised in Table 2 (the full set of measurements are provided in Table S1). Values of pH revealed that soils are either neutral or, mostly, alkaline (pH > 7.5), and exhibit little variation with transect distance or height or soil depth. Overall, pH is lowest at Sepidan (mean = 7.37) and greatest at Darab (7.91). Electrical conductivity (EC) is variable between and within the regions but is always non-saline (< 2 dS m^-1^) or very slightly saline (< 4 dS m^-1^). Values are mostly between 0.30 and 0.60 dS m^-1^ at Sepidan (mean = 0.43 dS m^-1^), with a small but persistent decrease with soil depth. By contrast, at Dasht Arjan, Darab and Sarvestan EC ranges from 0.35 to 2.27 dS m^-1^ (mean = 1.05 dS m^-1^), 0.58 to 1.92 dS m^-1^ (mean = 1.16 dS m^-1^) and 0.50 to 2.75 dS m^-1^ (mean = 1.50 dS m^-1^), respectively, and there are no systematic trends with soil depth.

**Table 2:** A summary of the physical and chemical properties of the soil samples by region, profile and horizon (horizon data are separated by /). EC = electrical conductivity; OC = organic carbon.

| Region | Profile | pH | EC, dS m<sup>-1</sup> | Sand, % | Clay, % | OC, % |
| --- | --- | --- | --- | --- | --- | --- |
| Dasht Arjan | DA1 | 7.66/7.75 | 0.56/1.05 | 33.7/36.9 | 17.5/21.0 | 1.73/0.23 |
|  | DA2 | 7.72/7.81/7.84 | 0.70/1.75/0.63 | 44.1/45.8/44.1 | 13.9/19.2/15.7 | 1.13/0.54/0.89 |
|  | DA3 | 7.89/7.48/7.85 | 1.66/1.40/2.27 | 22.7/24.4/22.7 | 21.0/22.8/26.4 | 2.14/2.63/1.03 |
|  | DA4 | 7.83/8.06/7.80 | 0.35/0.61/1.22 | 24.4/33.4/36.9 | 24.6/21.0/19.2 | 1.26/0.21/0.15 |
|  | DA5 | 7.98/7.45/7.92 | 0.61/0.52/0.47 | 38.7/49.4/51.2 | 19.2/19.2/15.7 | 0.83/0.19/0.21 |
|  | DA6 | 7.91/8.08/8.02 | 0.79/2.62/0.65 | 31.6/28.5/32.1 | 21.0/28.1/31.7 | 1.01/0.44/0.42 |
| Darab | DB1 | 7.80/7.90 | 0.89/0.86 | 28.0/28.0 | 22.8/33.5 | 1.03/0.64 |
|  | DB2 | 7.46/8.20/7.88 | 0.79/1.05/1.05 | 26.2/26.2/26.2 | 21.0/22.8/22.8 | 0.09/0.25/0.35 |
|  | DB3 | 7.63/7.75/7.93 | 1.66/1.92/1.48 | 38.4/20.3/16.3 | 23.9/38.0/38.0 | 1.06/0.54/0.45 |
|  | DB4 | 8.05/8.69 | 1.22/1.31 | 29.8/24.4 | 28.1/33.5 | 0.52/0.44 |
|  | DB5 | 7.60/7.96/7.96 | 0.50/1.40/1.13 | 40.5/29.8/28.0 | 15.7/31.7/29.9 | 0.54/0.15/0.35 |
|  | DB6 | 7.95/7.90 | 1.40/0.67 | 60.1/54.7 | 6.8/6.8 | 0.05/0.05 |
| Sarvestan | SA1 | 7.17/7.49 | 1.01/0.58 | 41.9/29.7 | 27.5/39.7 | 1.29/0.37 |
|  | SA2 | 7.63/7.57 | 0.69/0.53 | 41.7/29.8 | 16.1/31.7 | 1.08/0.37 |
|  | SA3 | 7.43/7.45 | 0.50/1.11 | 39.6/31.8 | 23.9/33.7 | 0.70/0.01 |
|  | SA4 | 7.52/7.49/7.37 | 0.52/1.17/2.08 | 37.8/23.8/33.8 | 23.7/39.6/31.8 | 0.37/0.05/0.29 |
|  | SA5 | 7.52/7.57/7.48 | 0.90/1.56/2.24 | 47.7/51.8/47.8 | 17.8/13.7/15.6 | 0.80/0.01/0.11 |
|  | SA6 | 7.51/7.55/7.55 | 1.79/2.24/2.25 | 45.9/55.9/58.1 | 13.7/3.5/1.5 | 0.98/0.13/0.27 |
|  | SA7 | 7.58/7.54/7.59/7.60 | 1.77/2.75/2.55/2.25 | 51.7/58.1/66.0/68.0 | 6.5/1.4/1.9/1.5 | 0.64/0.02/0.01/0.01 |
| Sepidan | SE1 | 7.30/7.37/7.36 | 0.47/0.43/0.40 | 23.0/19.5/23.0 | 27.8/36.7/36.7 | 2.53/0.58/0.58 |
|  | SE2 | 7.60/7.22/7.25 | 0.69/0.53/0.56 | 33.7/24.8/30.1 | 24.2/27.8/33.1 | 2.53/1.07/0.78 |
|  | SE3 | 7.34/7.37/7.39 | 0.47/0.39/0.41 | 24.8/21.2/28.4 | 27.8/36.7/33.1 | 1.65/0.87/0.87 |
|  | SE4 | 7.35/7.51/7.34 | 0.57/0.40/0.43 | 42.6/43.0/40.8 | 27.8/26.0/27.8 | 2.92/0.78/0.39 |
|  | SE5 | 7.28/7.30/7.40 | 0.46/0.35/0.33 | 33.7/30.1/26.6 | 22.4/27.8/34.9 | 1.17/0.39/0.39 |
|  | SE6 | 7.48/7.34/7.29 | 0.52/0.46/0.40 | 35.5/35.5/39.1 | 26.0/26.0/27.8 | 0.97/0.48/0.29 |
|  | SE7 | 7.39/7.32/7.37/7.45 | 0.40/0.30/0.32/0.26 | 31.9/23.0/30.1/35.5 | 26.0/27.8/31.4/29.6 | 0.48/0.19/0.25/0.05 |
|  | SE8 | 7.36/7.36/7.40 | 0.42/0.39/0.28 | 56.9/43.0/57.2 | 18.9/18.9/17.1 | 0.39/0.19/0.05 |

Organic carbon (OC) contents of the soils are variable between and within regions, with averages higher at Dasht Arjan and Sepidan (about 0.85%) than at Darab and Sarvestan (< 0.45%). Despite this variation, there is, in most cases, a reduction in OC with increasing soil depth. This reduction is most persistent at Sepidan and greatest, quantitatively, at Sarvestan, where values decrease by more than an order of magnitude in two profiles. Soil texture is summarised in terms of the percentages of sand and clay (with the remainder comprising silt). Thus, at Darab an increase in clay content and decrease in sand content was observed with increasing depth, while at Sepidan an increase in clay but similar sand content was generally found. At Dasht Arjan, there was a similar distribution of clay and sand with depth but at Sarvestan clay and sand distributions were more variable and a shift towards more sand and less clay was observed with increasing elevation along the transect.

#### 3.2. Microplastic abundance and distributions

MPs isolated from the soil samples are exemplified in Figure 2. In total, 392 MPs were retrieved, comprising 342 fibres, 48 fragments or (mainly) films, and two spherules. Overall, MPs were present in all but five soil samples and fibres were present in all but seven samples. Fibres appeared to be roughly equally distributed throughout each horizon at the four locations, and were most abundant at Sepidan (134 in 25 soils) and least abundant at Sarvestan (45 in 19 soils). Other shapes were least abundant at Sarvestan (1 in 19 soils) but most abundant at Dasht Arjan (24 in 17 soils). As a consequence, the ratio of other shapes to fibres ranged from about 0.02 at Sarvestan to about 0.36 at Dasht Arjan. Regarding individual profiles, the highest numbers of fibres (≥ 20) were observed in DB2, DB3 and SA7 (barren land), DB3 and SE1 (pasture), and SE7 (rainfed agriculture). The highest number of other shapes was encountered at three sites in Dasht Arjan consisting of agriculture, pasture and forest.

**Figure 2.**
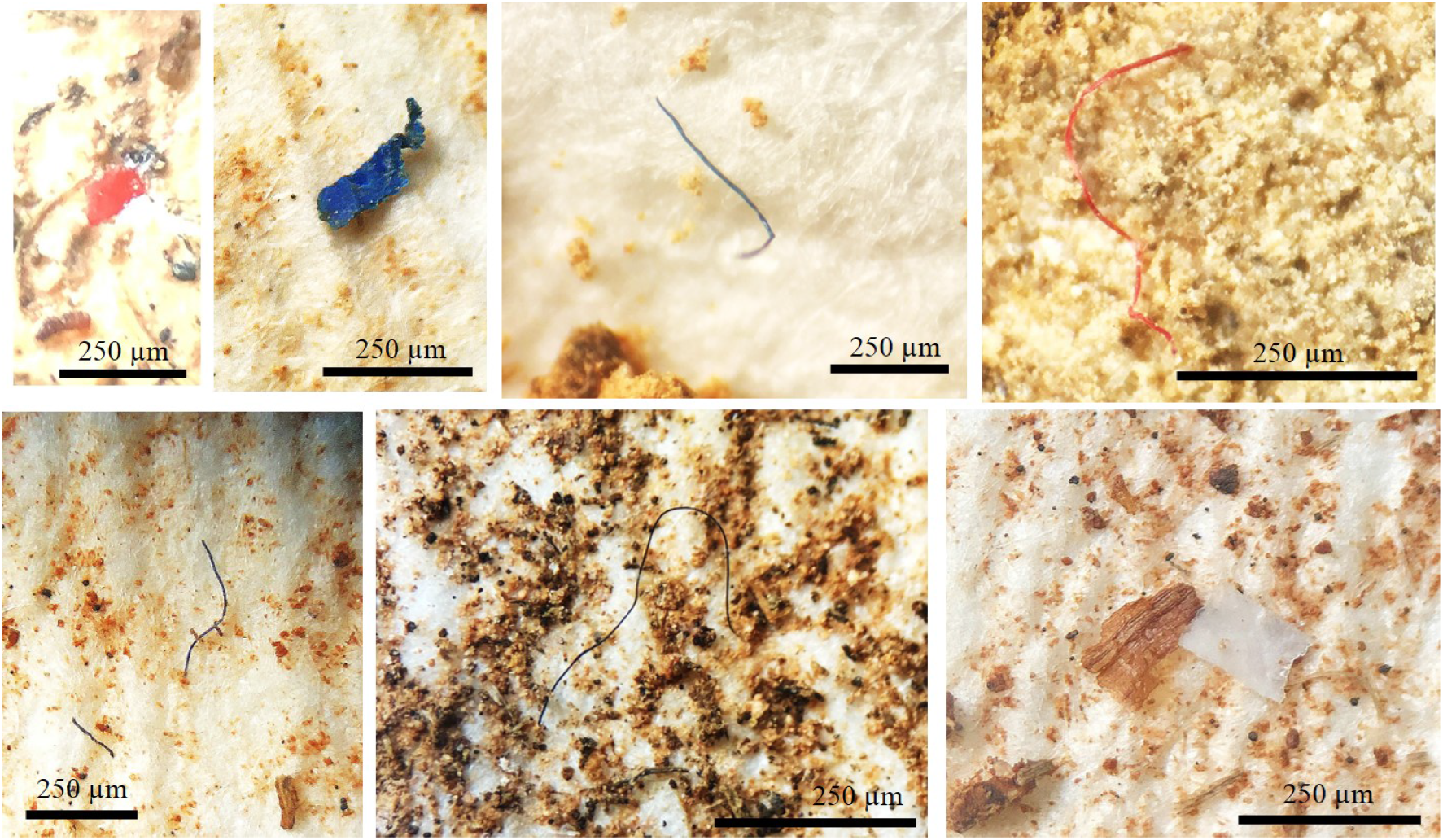
Examples of some of the fibrous and sheet-like MPs isolated from the soils in the present study and identified under the microscope.

With respect to MP size, 118 were small (< 100 μm) and 24 were large (> 500 μm), with ratios of small to large ranging from 2 at Sarvestan to about 7.2 at Darab. Fibres exhibited a range of colours but were mainly black or blue throughout, and while fragments and films displayed various colours, we noted mainly black or white at Darab, Sepidan and Sarvestan but a dominance of orange or pink/red at Dasht Arjan.

Correlation analysis revealed very few significant relationships between MP abundance and soil characteristics. Specifically, at Darab, where a downward decrease in the ratio of sand to clay was observed, MPs were inversely correlated with the percentage of sand (*r*_s_ = -0.551, *p* = 0.033; *n* = 15) and positively correlated with soluble Na content (*r*_s_ = 0.592, *p* = 0.020; *n* = 15), while at Sarvestan, where the ratio of sand to clay increased with increasing elevation, MPs were inversely correlated with the percentage of silt (*r*_s_ = -0.563, *p* = 0.012; *n* = 19) and positively correlated with soluble K content (*r*_s_ = 0.465, *p* = 0.025; *n* = 19).

Figure 3 shows the mean concentrations of MPs in the soil samples, calculated from the summed number of particles in each horizon normalised to 1 kg and divided by the number of samples per profile, by land use type. The highest mean concentration is where only one profile was sampled, with errors in the remaining cases highlighting the variation within land use type. Because of this variation, no significant differences in MP concentrations were observed amongst land use categories where *n* ≥ 2 (according to a Kruskal-Wallis test; *p* > 0.05).

**Figure 3.**
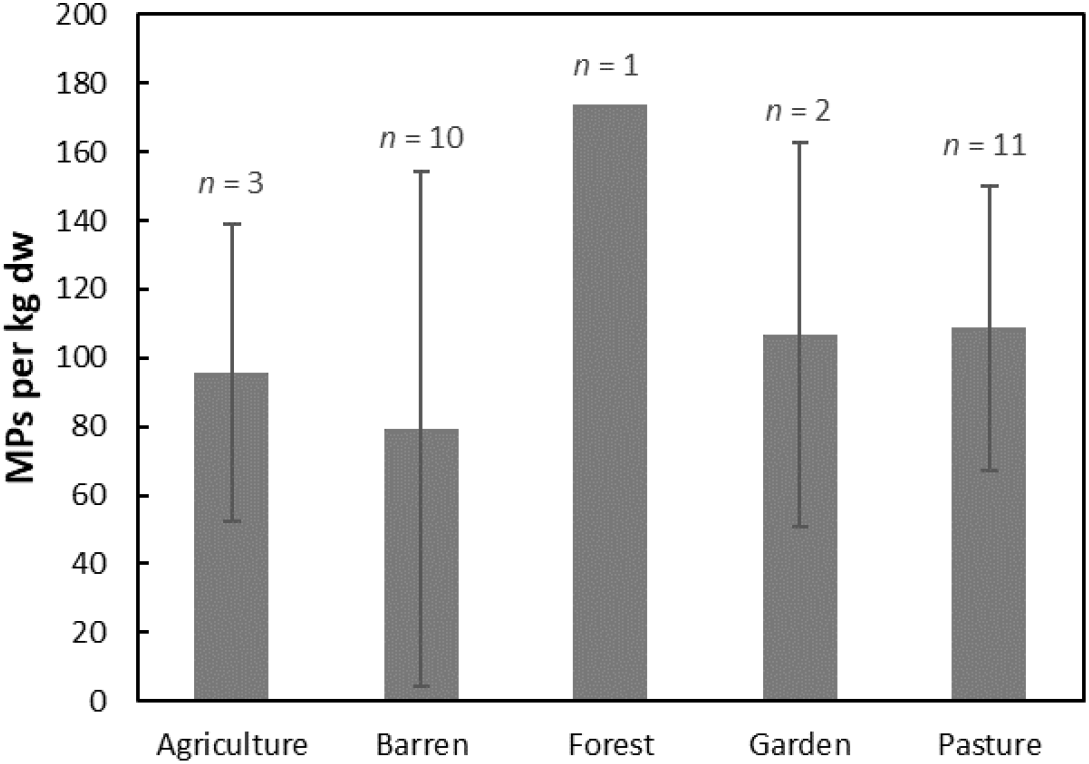
Mean and standard deviation (where errors are shown) of MP concentrations per kg of dry soil sample calculated from the summed number of particles per profile divided by the number of soils sampled in each profile (*n* is the number of profiles).

#### 3.3. Polymeric makeup of MPs

The results of μ-Raman analysis of MPs from eight profiles encompassing all four regions are summarised in Table 4. Polyamide, and mainly nylons, was the most widely distributed and abundant, although in a few cases nylon appeared to be blended with polypropylene. Polyester (including polyethylene terephthalate) and (pure) polypropylene were next in abundance but exhibited an unequal distribution between the regions (for example, the latter was absent from Dasht Arjan and the former absent from Sepidan). Polyethylene was detected in each region and polystyrene was absent from Sarvestan. Among the “other” category were acrylonitrile butadiene styrene, ethylene-vinyl acetate, various polydienes and the biopolymer, polylactic acid. Aside from the differences among the regions referred to above, polymers appeared to be heterogeneously distributed, with no evidence of fractionation by, for example, density, along transects or with profile depth.

**Table 3:** Number and size classification of microplastics (categorised as fibres and other) in 50-g (dry) soil samples by region, profile and horizon (horizon data are separated by /).

| Region | Profile | Fibres | Other | <100 µm | >500 µm |
| --- | --- | --- | --- | --- | --- |
| Dasht Arjan | DA1 | 2/3 | 0/0 | 1/0 | 0/0 |
|  | DA2 | 1/1/5 | 0/1/6 | 0/1/5 | 0/0/0 |
|  | DA3 | 6/0/5 | 0/1/0 | 4/1/0 | 2/0/1 |
|  | DA4 | 5/5/4 | 0/0/1 | 1/2/4 | 2/0/0 |
|  | DA5 | 3/6/2 | 0/3/5 | 2/5/5 | 0/0/0 |
|  | DA6 | 10/6/3 | 1/1/5 | 1/0/4 | 0/1/0 |
|  | total | 27/21/19 = 67 | 1/6/17 = 24 | 9/9/18 = 36 | 4/1/1 = 6 |
| Darab | DB1 | 2/5 | 3/3 | 4/3 | 0/0 |
|  | DB2 | 14/9/11 | 0/1/0 | 2/5/2 | 1/1/0 |
|  | DB3 | 2/16/8 | 0/0/0 | 0/4/2 | 0/1/1 |
|  | DB4 | 2/2 | 0/0 | 0/0 | 0/0 |
|  | DB5 | 5/9/6 | 0/0/1 | 3/3/3 | 0/1/0 |
|  | DB6 | 3/2 | 1/2 | 1/4 | 0/0 |
|  | total | 28/43/25 = 96 | 4/6/1 = 11 | 10/19/7 = 36 | 1/3/1 = 5 |
| Sarvestan | SA1 | 4/3 | 0/0 | 0/2 | 0/0 |
|  | SA2 | 0/0 | 0/0 | 0/0 | 0/0 |
|  | SA3 | 1/3 | 0/0 | 0/0 | 1/0 |
|  | SA4 | 1/0/1 | 0/0/0 | 0/0 | 0/0 |
|  | SA5 | 4/1/2 | 0/1/0 | 0/0/1 | 1/2/0 |
|  | SA6 | 0/1/0 | 0/0/0 | 0/0/0 | 0/0/0 |
|  | SA7 | 4/3/7/10 | 0/0/0/0 | 2/0/5/2 | 1/0/0/1 |
|  | total | 14/11/10/10 = 45 | 0/1/0/0 = 1 | 2/2/6/2 = 12 | 3/2/0/1 = 6 |
| Sepidan | SE1 | 7/5/10 | 0/0/1 | 2/1/1 | 1/1/0 |
|  | SE2 | 8/4/4 | 3/2/1 | 1/2/0 | 1/0/0 |
|  | SE3 | 10/3/6 | 2/0/2 | 5/0/3 | 1/0/0 |
|  | SE4 | 2/1/4 | 1/0/0 | 1/0/1 | 0/0/0 |
|  | SE5 | 8/6/1 | 0/0/0 | 1/2/0 | 0/0/1 |
|  | SE6 | 2/0/7 | 0/1/0 | 0/1/3 | 0/0/0 |
|  | SE7 | 3/9/6/9 | 0/0/0/1 | 0/3/2/3 | 0/0/0/0 |
|  | SE8 | 8/5/6 | 0/0/0 | 1/0/1 | 1/1/0 |
|  | total | 48/33/44/9 = 134 | 6/3/4/1 = 14 | 11/9/11/3 = 34 | 4/2/1/0 = 7 |

**Table 4:** Distribution of polymers in 98 MPs isolated from soil samples across the four regions.

| Polymer | Dasht Arjan | Darab | Sepidan | Sarvestan | Total |
| --- | --- | --- | --- | --- | --- |
| Polyamide | 5 | 24 | 12 | 4 | 45 |
| Polyester | 7 | 3 | 0 | 3 | 13 |
| Polyethylene | 2 | 3 | 2 | 1 | 8 |
| Polypropylene | 0 | 6 | 6 | 1 | 13 |
| Polystyrene | 1 | 2 | 1 | 0 | 4 |
| Other or unidentified | 1 | 10<sup>a</sup> | 4 | 0 | 15 |
| Total | 16 | 48 | 25 | 9 | 98 |

<sup>a</sup> Includes one sample that was inorganic.

### 4. Discussion

The results of the study reveal that MPs are ubiquitous throughout soils at concentrations up to about 200 per kg but are heterogeneously distributed in terms of size, shape, colour and polymer, and with respect to location, land use, horizon and soil characteristics. Results reported in the literature where MPs have been determined in surficial or rhizospheric soils encompassing different land uses are generally heterogeneous, but more distinct differences between land uses or relationships with environmental variables are often observed, albeit not always with statistical confirmation. For instance, differences in soil MP abundance have been reported among different ecosystems (Álvarez-Lopeztello et al., 2021; Xu et al., 2022), different agricultural practices (Liu et al., 2022; Zhang et al., 2022a; Bi et al., 2023; He et al., 2023), and different agricultural practices, urban areas and woodland (Qiu et al., 2023; Yoon et al., 2024), and between managed and unmanaged land (Corradini et al., 2021) and agricultural land and urban areas (Akca et al., 2024). By contrast, and consistent with our study, Zhang et al. (2024) report no differences in soil MP abundance between industrial land, different agricultural fields and woodland. Relationships (or trends) have been reported between MP abundance and soil properties, including clay-sand content, pH, bulk density, organic matter and nitrogen (Álvarez-Lopeztello et al., 2021; Liu et al., 2022; Zhang et al., 2022b; Qiu et al., 2023; He et al., 2023; Zhang et al., 2024), climatic factors, such as precipitation, surface temperature and wind speed (Zhang et al., 2022a), and population density (Bi et al., 2023). Moreover, we observed significant relationships with soluble Na or K content and the inverse of sand or silt at two locations. However, it could be argued that these relationships are not necessarily related to land use but are some function of more general environmental or anthropogenic variables.

In a review of MP abundance in agricultural soils of China, where most studies above have been undertaken, Liu et al. (2023) suggest that mulching using plastic films is responsible for the variations observed, at least in the agricultural sector. Accordingly, contamination depends on the type of plastic used, the period and length of mulching and the degree of weathering effected by, for example, irrigation. Mulching cannot, however, account for the results observed here, where the practice is not common and the most abundant MPs were persistently fibres (rather than films) that extended beyond cultivated land (well water- and rainfed-irrigated gardens and agriculture) into forest and barren land. Fibres have relatively low settling velocities and high degrees of horizontal and upward motion in air, allowing them to be transported long distances and become heterogeneously dispersed (Abbasi et al., 2023; Preston et al., 2023; Xiao et al., 2023). Subsequent deposition may be dry or wet, with the majority of studies concluding that washout with rainfall is more important either directly (Szewc et al., 2021; Kyriakoudes and Turner, 2023; Yuan et al., 2023) or, under vegetation, via combing-out (filtering) of particles by the canopy and washing off during precipitation (Klein and Fischer, 2019). Dry and wet deposition, combing-out and rainfed agricultural practices are all likely to contribute to contamination by fibrous MPs in the region under study in Iran.

The present investigation has also shown the propensity for MPs to migrate to depths beyond 1 m in soils of various land uses. Previous studies have found MPs below the soil surface, although most report arbitrary depths of a few tens of cm and not distinctive soil horizons. For example, Zhang and Liu (2018) found similar MP concentrations (and mainly fibres) in surface (0 to 5 cm) and subsurface (5 to 10 cm) agricultural soils, while Xu et al. (2022) observed higher MP concentrations in subsurface (10 to 20 cm) soils than surface (0 to 10 cm) soils of artificial ecosystems (plantations) but higher concentrations below the surface in natural ecosystems (forests). Yu et al. (2021) found that smaller MPs (< 0.5 mm), and mainly fragments and films, tended to migrate to deeper layers (10 to 25 cm) in various agricultural soils. Li et al. (2023) examined MPs (of various shapes) in three soil depths down to 60 cm of seven land use types (mainly different agricultural practices). MPs of various shapes were encountered throughout the soil profiles but both abundance and size tended to decline with depth.

There are a number of mechanisms by which MPs deposited at the soil surface may penetrate deeper horizons. Being a porous medium, soil allows the migration (or leaching) of particles with percolating precipitation under gravity. In theory, smaller particles (including those generated by weathering in situ) and those that are denser and more hydrophobic are more mobile, while longer fibres are subject to greater retardation (Engdahl, 2018; Gao et al., 2021). Our observations do not indicate a shift in MP size, shape or density-hydrophobicity (polymer type) with depth or soil particle size, however, requiring the presence of additional vertical transport mechanisms. Where land is managed, disturbance can result in homogenisation to the tilling depth, although typically this is down to about 25 cm (Hobson et al., 2022). Bioturbation by earthworms (through adhesion, ingestion-egestion or burrowing) and plant roots (through movement, expansion, water retention or decomposition) may facilitate downward migration to 50 cm or more (Gabet et al., 2003; Rillig et al., 2017; Li et al., 2020), as might soil cracking in dry climates. Column experiments have shown that granular and spherical MPs are able to penetrate to greater depths in soils over successive wet-dry cycles, with calculations suggesting penetration depths below 5 m over a 100-year period are possible (O’Connor et al., 2019). The extents to which longer but more flexible and irregular fibres are able to migrate by this process are, however, unknown.

The significance or relevance of each mechanism above is unclear in the present context. However, the similarities in MP shape, size, colour and polymer throughout the soil depths sampled and across the region studied suggest that any vertical (or horizontal) sorting is not specific to these characteristics and that any in situ physico-chemical or biological degradation into smaller particles is not discernible. Moreover, although we do not know the precise timescales involved in vertical migration in the region under study (we may infer it might be on the order of a decade based on the calculations by O’Connor et al. (2019)), our results suggest that, quantitatively and qualitatively, MP input at the soil surface has been rather consistent over this period. The depth to which we have observed MP migration also suggests that subterranean ecosystems and ground water may be readily contaminated. Moreover, given the ubiquity of MP deposition from the atmosphere, contamination of these resources is not necessarily specific to locations where point sources of MPs exist.

## 5. Conclusions

This study has shown that MPs, dominated by dark-coloured fibres and up to concentrations of about 200 kg^-1^, are ubiquitous in soils from four regions of Fars Province, Iran, that encompass various land uses, and extend to the deepest horizons excavated (140 cm). Distributions are, however, heterogeneous in terms of abundance, size and polymeric make up and with respect to land use, elevation, soil depth and soil characteristics. With few or no direct inputs from point sources related to agriculture, urbanisation or industry, for example, the main vehicle for accumulation appears to be atmospheric deposition at the soil surface, with subsequent downward migration likely assisted by precipitation, bioturbation and cracking of the soil structure through wetting-drying cycles. Migration poses risks to subterranean ecosystems and impacts on groundwater quality that are currently poorly defined.

## Acknowledgements

We gratefully acknowledge Shiraz University for funding this Ph.D. thesis (Grant Number: 2GRC1M371631) and the Centre for Environmental Studies and Emerging Pollutants (ZISTANO) for their technical support.

## Data Availability

Data will be available upon request from Sajjad Abbasi.

## Notes

### Competing Interest Statement

The authors have declared no competing interest.

